# Visual Biochemistry: modular microfluidics enables kinetic insight from time-resolved cryo-EM

**DOI:** 10.1101/2020.03.04.972604

**Authors:** Märt-Erik Mäeots, Byungjin Lee, Andrea Nans, Seung-Geun Jeong, Mohammad M. N. Esfahani, Daniel J. Smith, Chang-Soo Lee, Sung Sik Lee, Matthias Peter, Radoslav I. Enchev

## Abstract

Mechanistic understanding of biochemical reactions requires structural and kinetic characterization of the underlying chemical processes. However, no single experimental technique can provide this information in a broadly applicable manner and thus structural studies of static macromolecules are often complemented by biophysical analysis. Moreover, the common strategy of utilizing mutants or crosslinking probes to stabilize otherwise short-lived reaction intermediates is prone to trapping off-pathway artefacts and precludes determining the order of molecular events. To overcome these limitations and allow visualisation of biochemical processes at near-atomic spatial resolution and millisecond time scales, we developed a time-resolved sample preparation method for cryo-electron microscopy (trEM). We integrated a modular microfluidic device, featuring a 3D-mixing unit and a delay line of variable length, with a gas-assisted nozzle and motorised plunge-freeze set-up that enables automated, fast, and blot-free sample vitrification. This sample preparation not only preserves high-resolution structural detail but also substantially improves protein distribution across the vitreous ice. We validated the method by examining the formation of RecA filaments on single-stranded DNA. We could reliably visualise reaction intermediates of early filament growth across three orders of magnitude on sub-second timescales. Quantification of the trEM data allowed us to characterize the kinetics of RecA filament growth. The trEM method reported here is versatile, easy to reproduce and thus readily adaptable to a broad spectrum of fundamental questions in biology.

## Introduction

The structures of biological macromolecules largely determine biochemical activity, mechanism, and specificity through highly dynamic conformational changes. This remodelling is coupled to functions such as enzyme catalysis, allostery, and interactions with other macromolecules, typically occurring on microsecond to millisecond timescales^1,2^. Such structural rearrangements proceed through energetically accessible transition-states^3^ without destabilizing the overall fold necessary for function^4^. Early studies demonstrated that proteins exist in a variety of conformational sub-states that correspond to local energy minima^5^. However, it is experimentally challenging to isolate these states for structural studies and relate them to functionally relevant intermediates.

Cryo-electron microscopy (cryo-EM) and single-particle analysis are widely used to determine structures of biological macromolecules^6^. One distinct advantage of the method is that the samples are prepared in solution under nearly native conditions, allowing the imaging of numerous instances of individual specimens and application of statistical methods to disentangle compositional and conformational variability. Indeed, recent studies have shown that many separate states can be identified and solved to high resolution within a cryo-EM sample^4,7^. However, such workflows do not directly identify functionally relevant states, nor do they elucidate the sequence in time of structural transitions underpinning a reaction pathway. Moreover, short-lived active intermediates, which are rare under equilibrium conditions, remain undetected. Therefore, there has been a long-standing interest in developing time-resolved cryo-EM sample preparation and analysis to visualise such states. First successes came from cryo-electron crystallography^8–12^ by diffusing small interactants into crystals, broadly analogous to time-resolved X-ray crystallography techniques^13^ and limited to crystalline samples and small diffusible ligands. More recent attempts in the framework of cryo-EM and single-particle analysis have traced structural changes over time in processes that proceed over several seconds and even minutes. This was only possible because the studied processes were slower than the time required to prepare a cryo-EM sample grid by the standard method of manual application, followed by automated blotting and plunge-freezing^14–17^. Dynamic structural states could also be resolved by freeze-trapping samples, pre-incubated at different temperatures^18^. However, both of these techniques are applicable to a small subset of biochemical reactions that have slow kinetics or show substantial temperature sensitivity, and do not enrich short-lived intermediates.

To overcome these limitations and develop a general time-resolved sample preparation method for cryo-*EM* (trEM) requires building a miniaturized mixer and bioreactor able to rapidly initiate and synchronize biochemical reactions, followed by spreading the incubated sample onto a cryo-EM grid without the need for manual operation or blotting, collectively faster than the lifetime of the structures of interest^19^. Previous work has shown that combining reactants in microfluidic devices followed by rapid application of sample by gas-assisted spraying is in principle possible^20–25^, and has yielded fascinating new insights into biology^26^. However, substantial technical challenges remain unaddressed. For instance, in order to widen the applicability of the method, developments are needed to reduce the complexity of microfluidics design and manufacturing without diminishing the reliability and reproducibility of results. Additionally, gas-assisted aerosol generation for blot-free sample application through a microfluidic device has not been studied systematically. Lastly, the ability to achieve reliable mixing on millisecond timescales inside microfluidic devices has not been experimentally assessed in the context of trEM, nor have the errors on nominal in-chip incubation times been measured. Collectively these pose major impediments to conducting well-controlled time-resolved structural studies by cryo-EM, as have been reported by time-resolved x-ray crystallography and free electron laser experiments^27^.

Here we address most of the above by developing a trEM workflow that incorporates a modular microfluidic platform with an *in situ* 3D mixer and tunable incubation time from ten to thousands of milliseconds, capable of enriching intermediate states of biochemical reactions on cryo-EM grids. The system is largely automated and simple to manufacture without access to specialised facilities. Comprehensive assessment of the obtained cryo-EM sample quality demonstrated the unique advantages of rapid blot-free sample vitrification. Moreover, we established a new biochemical model system, derived from the bacterial homologous recombination pathway, which allowed us to characterise the obtainable time-resolution in quantifiable terms. We demonstrate the ability to extract kinetic information from structural changes on timescales spanning three orders of magnitude pertinent to macromolecular function. Combining all the above developments we present a robust methodology for the study of dynamic structures by trEM.

## Results

### An integrated system to allow time-resolved preparation of cryo-EM samples

In standard cryo-EM sample preparation, several microliters of sample are pipetted onto a support grid, followed by removing most of the sample liquid with blotting paper (**Figure 1A**). This leaves a very thin layer of sample that can be vitrified under ambient pressure and imaged with high contrast and minimal secondary scattering^28^. However, blotting requires at least a few seconds and is thus incompatible with time-resolved studies of most biochemical processes. To prepare a cryo-EM sample on millisecond timescales, we developed an integrated sample preparation method with substantial automation (**Figure 1B**). This system (**Supp Figure 1A**) utilises microfluidic devices that rapidly and reliably mix and then incubate reactants on timescales relevant to the studied biochemical process. A gas-assisted nozzle generates a sample aerosol, which is applied, blot-free, onto a standard cryo-EM support grid and rapidly plunged into liquid ethane by a servo motor. The assembly is modular so that different microfluidic devices can be operated by the same set-up, ensuring high reproducibility. Additionally, the system can be placed into an environmental chamber with humidity and temperature control. An integrated electronic board and software were developed to coordinate the function of all components (**Supp Figure 1B, C**). This control system is instrumental to reduce sample volume, as it ensures constant flow for a minimal time before spraying the sample onto the grid. The design principles and optimised working specifications of each major component are described in detail below.

**Figure 1.**
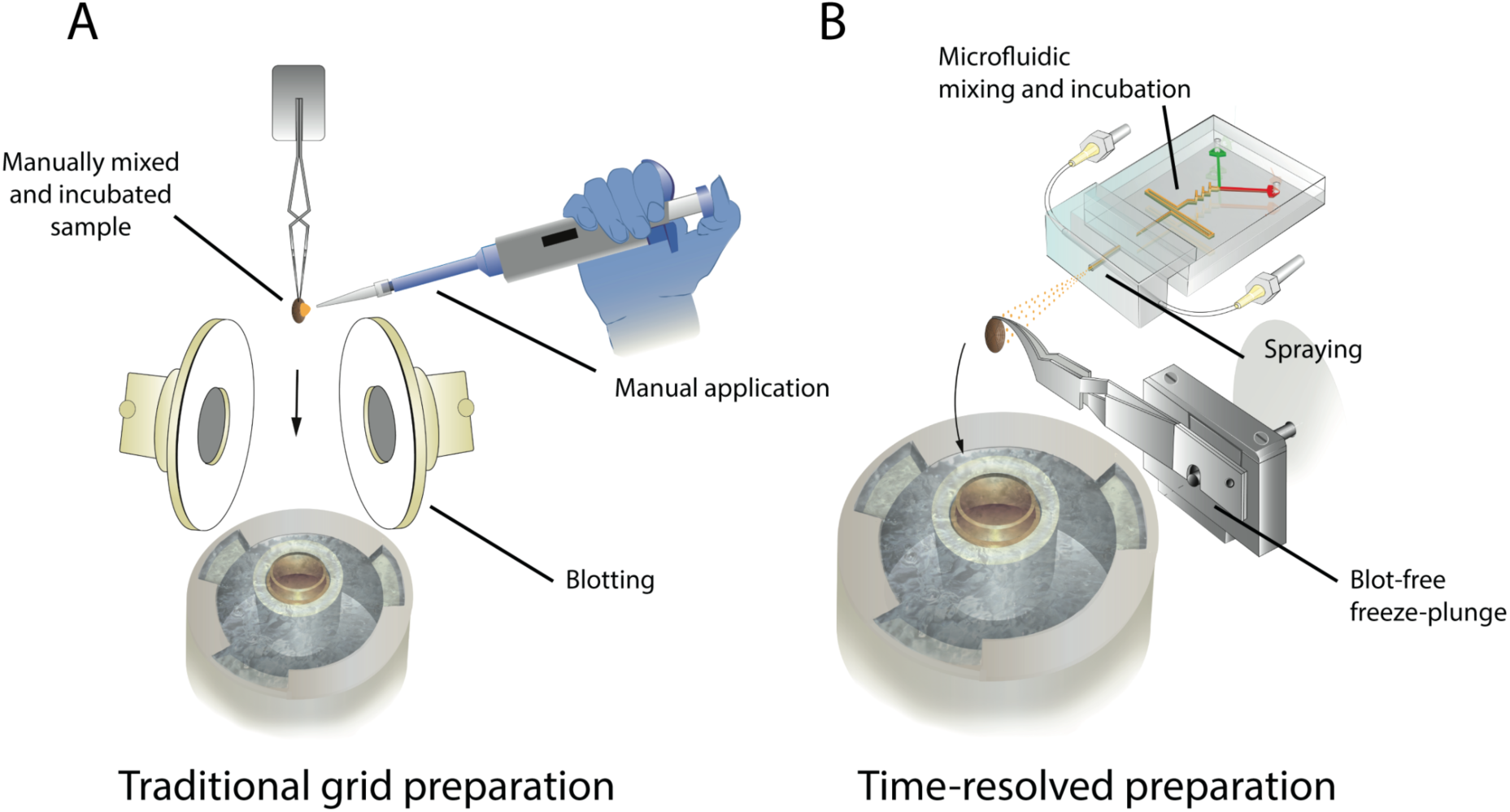
Sample preparation methods for cryo-EM. **A** Drawing of the standard method, which entails manually pipetting several microliters of sample onto a support grid, removing the excess liquid by blotting (touching with filter paper) and vitrification by plunge-freezing into liquid ethane. The most popular commercial system for this preparation is marketed as Vitrobot, and this name is used interchangeably with standard cryo-EM sample preparation throughout the text. **B** Drawing of time-resolved cryo-EM sample preparation. A microfluidic device mixes and incubates a biochemical reaction, then sprays the sample onto a plunging grid such that blotting is not required, and finally the sample grid is plunged into liquid ethane to achieve vitrification.

### Rapid microfluidic mixing to initiate biochemical reactions

The first critical element is the initiation of the biochemical reaction inside the microfluidic chip by mixing. Kinetic rates that determine the formation and lifetimes of reaction intermediates are dependent on the local concentration of the components^29,30^. Thus, to reliably enrich for a specific state at the endpoint of freezing, the starting point must be precisely defined by optimizing the mixing of the individual reactants. We adopted an *in situ* three-dimensional (3D) mixer, previously reported to significantly enhance mixing efficiency^31^ **(Figure 2A, Supp Figure 2A)**. In the mixing region, the bulk fluid flow is along the axis of the channel but fluid around a channel bend has three-dimensional flow combined with secondary flows, generated in the channel cross-section. These secondary flows integrated with the axial flow distort and stretch interfaces and can produce chaotic advection. Thus, the interfacial area across which diffusion occurs is greatly increased, and enables near-perfect mixing in millisecond time and nanolitre volumes.

**Figure 2.**
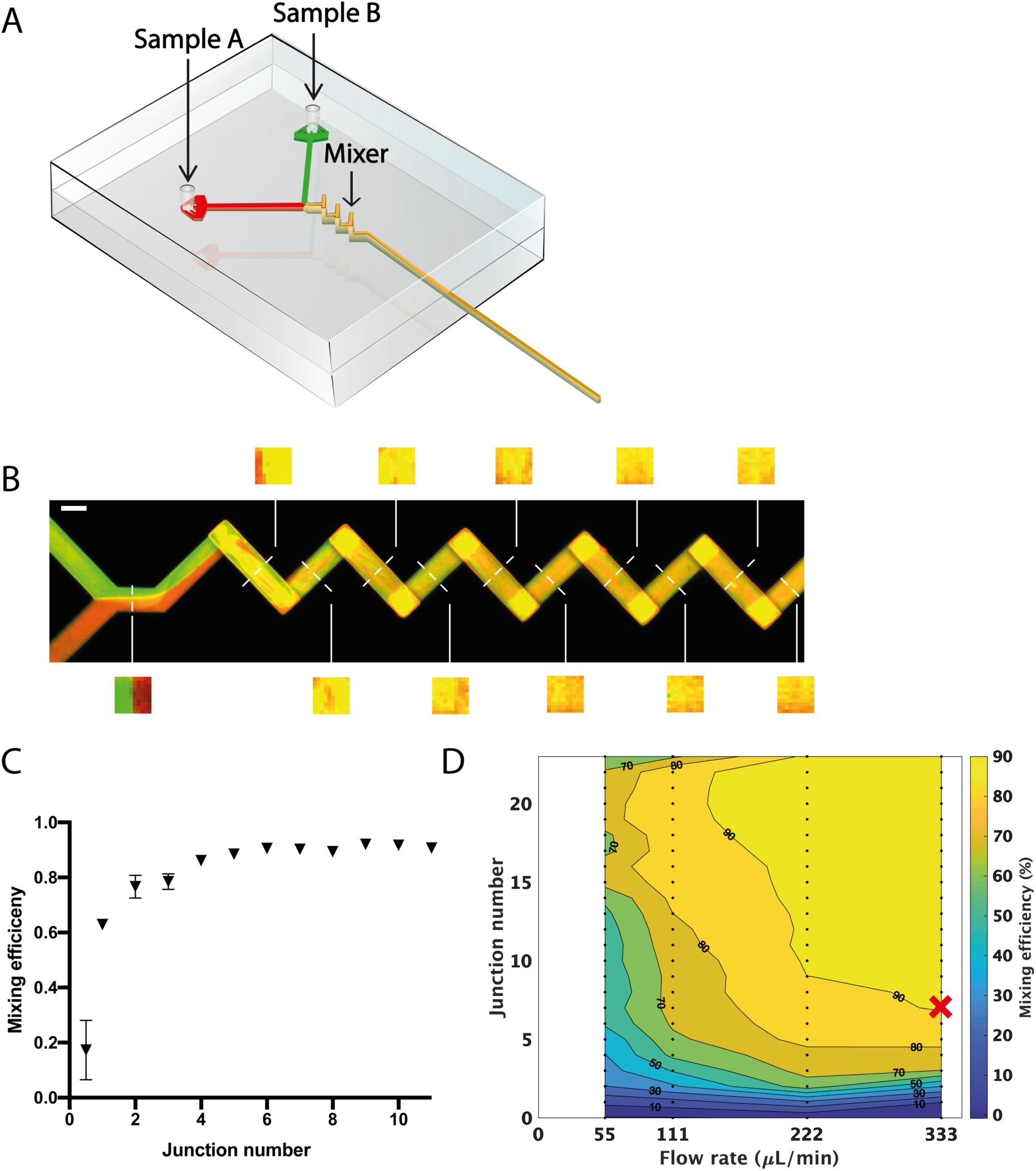
Rapid and reliable sample mixing in a microfluidic device. **A** Schematic representaron of a microfluidic device with the mixer geometry shown in yellow. **B** Confocal micrograph representing an optical slice through the centre of a mixing microfluidic geometry at steady state flow, 333 μl/min per channel, of two fluorescent dyes. Transversal slice reconstructions through the channel used for quantification of the mixing efficiency are shown. Scale bar is 100 μm. **C** Quantification of mixing efficiency data shown in **B** as a function of 3D mixing elements (junctions). **D** Phase diagram representaron of mixing efficiency as a function of liquid flow rate (per channel) and number of 3D mixing elements. The condition minimising both flow rate and mixing channel length used in the remainder of this work is indicated (red cross).

To validate the above theoretical consideration and to quantify the mixing efficiency and time, we developed a dedicated experimental workflow. We produced a microfluidic device composed of a large number of sequential 3D-mixing elements (junctions), pumped two distinguishable fluorophore solutions through the two channels and mounted it on a confocal microscope to image the mixing along the channel as a function of 3D junctions and flow rates (**Figure 2B, C, Supp Figure 2B, C**). The mixing efficiency and time were then determined by image analysis of cross-sections along the channel (**Figure 2B, C**). We observed a secondary flow-induced thinning striation across the channel with increasing number of junction and flow rates, resulting from a vigorous mixing of two discrete flows **(Figure 2B, Supp Figure 2B)**. The sample flow rate and size of the mixing region were chosen to minimize the mixing time and required flow rate (**Figure 2D, Supp Figure 2C**). Based on this analysis, we chose a flow rate of 333 μl/min per channel and 7 mixing junctions for all experiments reported below, as it reached maximum mixing efficiency observed in the shortest amount of junctions.

### Rapid, blot-free cryo-EM sample preparation

After initiating the reaction inside the microfluidic device, the sample needs to be rapidly delivered to a cryo-EM support grid without compromising reaction synchronisation. To achieve this, we developed a gas-assisted nozzle (**Supp Figure 3A**) that produces a fine sample aerosol applicable without the need for time-consuming blotting. The nozzle was tailored to the microfluidic geometry and liquid flow rate determined above, with sample flowing at 666 μl/min out of a 100-micrometre circular orifice. A concentric gas stream breaks the sample jet into small droplets and accelerates them towards the grid, which allows them to spread on impact into a thin enough film for cryo-EM imaging. The nozzle assembles modularly with the microfluidic chips (**Figure 3A**), which allows integrating devices with distinct properties. Increasing the gas pressure produces smaller, albeit more irregular droplets (**Figure 3B, C**) but also increased droplet velocity (**Figure 3D, Supp Movie 1, 2**). Nitrogen gas pressure of 0.8 bar was found to be optimal to generate small enough droplets to spread into thin layers on the grid without significant damage to the support grid.

**Figure 3.**
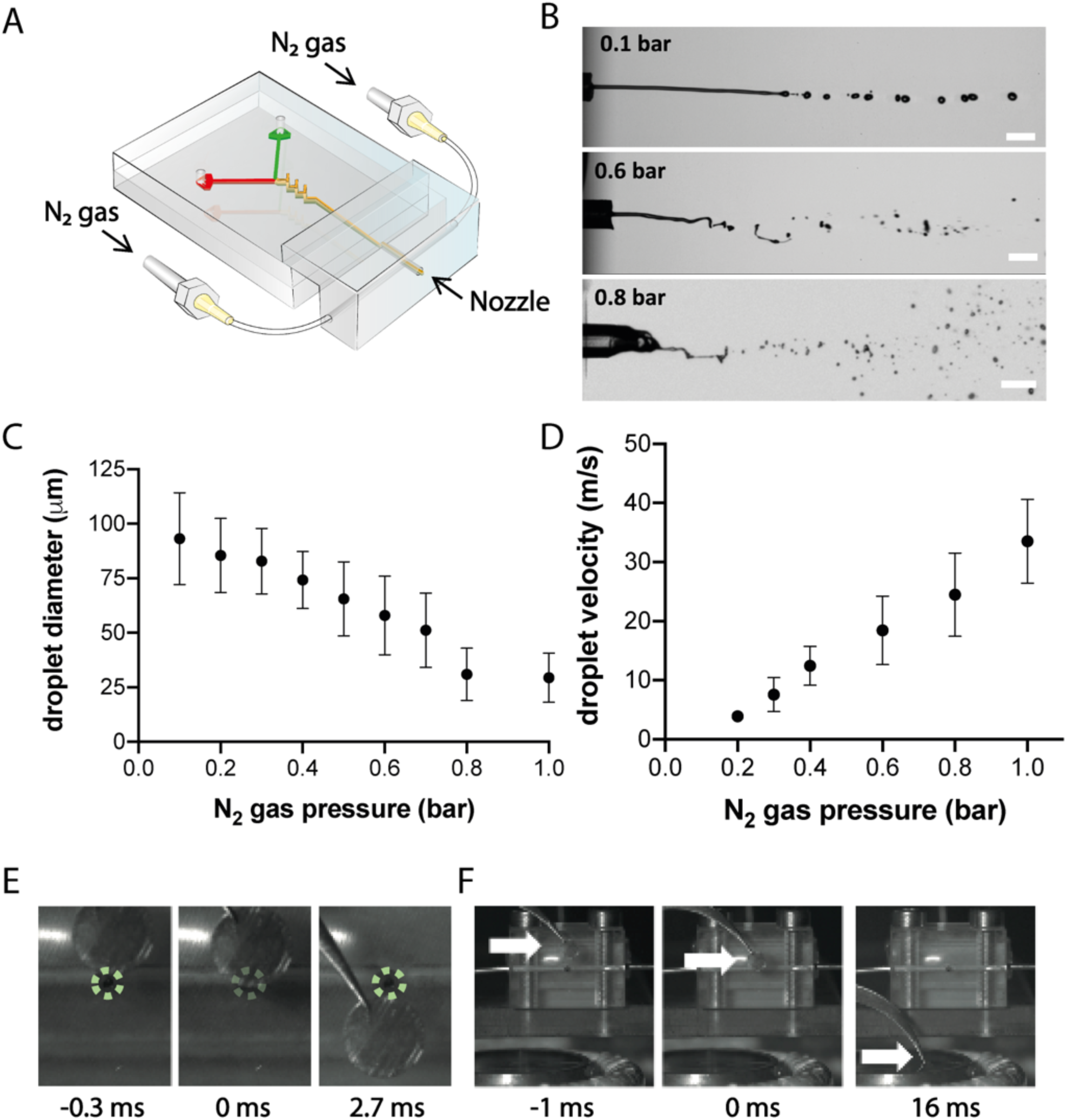
Sample spray and blot-free delivery onto a TEM support grid. **A** Schematic representaron of a microfluidic device with a gas-assisted spraying nozzle. **B** Snapshots from high-speed camera datasets of sample sprayed at indicated nominal gas pressure. Scale bar is 363 μm. **C, D** Quantification of droplet diameters **(C)** and speed **(D)** as a function of increased gas pressure. **E, F** Snapshots from high-speed camera imaging of a cryo-EM sample grid plunging **(E)** past the nozzle (nozzle position indicated in dotted green) and **(F)** from the nozzle into the liquid ethane container (positions indicated by white arrows), with time stamps in milliseconds.

The next critical step was the collection of sample aerosol onto a grid and its subsequent vitrification. This was done by attaching a pair of tweezers to a servo motor and radially plunging the grid into a liquid ethane container at a tuneable speed (**Figure 1B, Supp Figure 1A**). Servo motors are affordable and easily customizable, for example by combining with gears to increase or decrease the plunging speed. We found that plunging at 100 rpm (∼1.1 m/s radial speed) at a distance of 8 mm from the nozzle consistently produced grids with a sufficient number of areas suitable for cryo-EM imaging and high resolution single-particle analysis. This results in an integration time of sample on the grid of 2.7 ms (**Figure 3E**), and 16 ms between mixing of the sample to vitrification (**Figure 3F**). Faster plunging was not found to be beneficial as it reduced the amount of sample collected on the grid, while as expected, slower plunging fails to vitrify the sample. These parameters define the preparation time to around 20 ms, along with an integration time of ∼3 ms. The time of flight of the sample spray and subsequent vitrification was less than 2 ms^32^. Together with the 3 ms required for sample mixing (see above), the resulting dead time is less than 30 ms, which is fast enough to approach many hitherto unstudied biological reaction timescales.

### Quality assessment of cryo-EM sample prepared by blot-free spray-plunging

We applied trEM sample preparation to a series of different commercially available sample support grids and biological samples to assess the overall quality of ice and amount of microscopically collectable areas (**Figure 4A-C**). At the optimised plunging speed and distance from the nozzle to grid described above, we routinely observed sufficient numbers of grid squares containing sample droplets with good contrast. We detected thinner spreading over holey Quantifoil grids, but carbon-coated commercially available Quantifoil and lacey grids covered with a thin carbon layer also produced useful electron micrographs (**Figure 4B, C**). The particles were observed to be unperturbed, albeit with a similar but slightly lower concentration of protein compared to the standard procedure, as also noted in other rapid sample application methods^33^. For the remainder of this work, we focus on the most widely used grid type for cryo-EM, holey Quantifoil.

**Figure 4.**
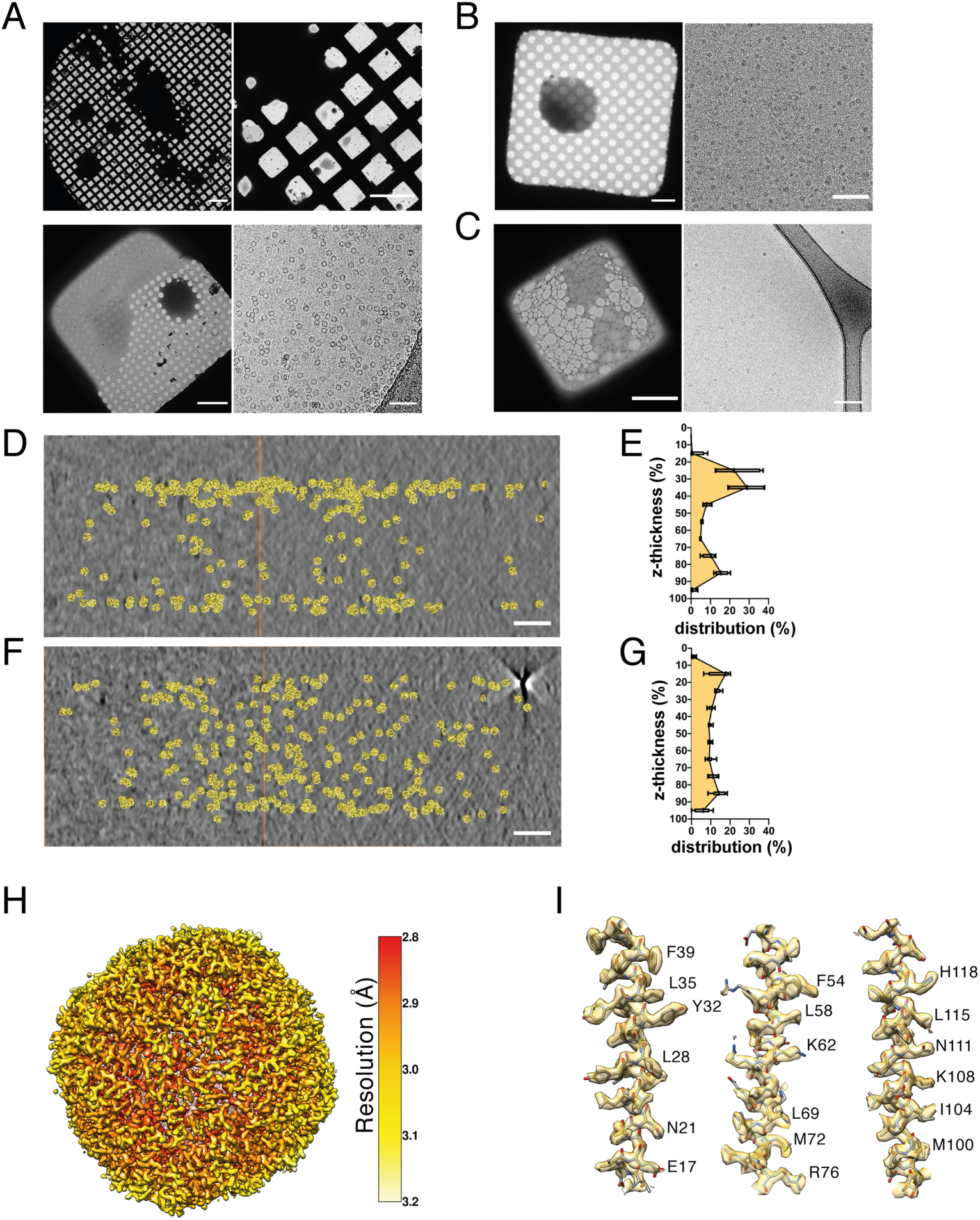
Assessment of sample cryo-EM samples prepared by blot-free spray-plunging. **A** Apoferritin sprayed on a holey Quantifoil grid. Scale bars represent 200 μm in upper left panel, 100 μm in upper right panel, 10 μm in lower left panel, 50 nm in lower right panel. **B** CSN^5H138A^-Cul1-N8/Rbx1 complex^79^ on a Quantifoil grid covered with thin carbon. Scale bars are 10 μm in left panel, and 100 nm in right panel. **C** 20S proteasome^80^ on a lacey grid covered with thin carbon. Scale bars are 5 μm in left panel, and 10 nm in right panel. **D-G** Tomographic reconstructions and segmentation of apoferritin distribution across the depth of a vitreous sample spanning a hole of a Quantifoil grid produced by the standard method **(D)** and blot-free spray-plunging **(F).** Scale bars are 50 nm. Quantification of the distribution of sample as a function of depth (in 10% steps) produced by conventional **(E)** and blot-free spray-plunging **(G)** from 15 independent measurements for each sample. **H** Single-particle reconstruction of apoferritin from a sample prepared by blot-free spray-plunging at global resolution of 2.7 Å, local resolution indicated by the given colour code. **I** Representative fit of atomic model and density of indicated regions.

To assess the sample quality obtained by this method we first performed tomographic reconstructions to determine the protein distribution relative to the air-water interfaces^34–36^. We used apoferritin as a model protein and compared biochemically identical samples prepared by trEM to the standard technique (**Figure 4D-G, Supp Figure 4A**). We obtained excellent contrast, although the average ice thickness was somewhat greater than 100 nm. To facilitate direct comparison, we collected tomograms from regions of matching ice thickness on grids prepared using a Vitrobot. Consistent with previous reports^36–38^ we observed a pronounced bi-phasic distribution of sample due to clustering at the air-water interfaces when prepared on a Vitrobot (**Figure 4D**) among multiple areas of independently produced grids (**Figure 4E**). Importantly, following the same analysis and quantifications of sample grids prepared by trEM we observed a significantly more uniform distribution of protein along the z-direction (**Figure 4F, G**), indicating an important improvement to sample quality.

To further characterise the quality of the sample, we collected datasets from both preparation methods for high-resolution structure determination by single-particle analysis. Datasets of similar size were collected and subjected to an identical single-particle analysis workflow (**Supp Figure 4B, C**). We were able to confirm that high-resolution structure of apoferritin could be reconstructed from data obtained by trEM grid preparation and that all expected near-atomic resolution details were retrieved (**Figure 4 H, I**). Importantly, the comparative analysis showed no significant difference between the datasets, with similar numbers of particles picked, similar proportions retained at each round of processing, and similar final resolution achieved (**Supp Figure 4B, C**). We thus conclude that the established trEM workflow reliably produces high-quality cryo-EM samples on sub-second timescales.

### Trapping pre-steady state kinetic intermediates by cryo-EM

To assess the fidelity of synchronization, and the ability to visualise pre-steady state reaction intermediates by trEM, we sought a biochemical test sample with broad significance to biology, and well-understood kinetics and steady-state structure, which would be readily correlatable in cryo-EM micrographs without recourse to novel image processing or quantification algorithms. We chose to study the prototypical bacterial recombinase RecA, which mediates homologous recombination pathways^39^ and is thus a critical regulator of DNA damage repair and antibiotic resistance arising from horizontal gene transfer. In particular, we probed the known kinetic behaviour of ATPγS-dependent growth of *E. coli* RecA protein on single-stranded DNA. RecA-ssDNA helical filaments are simple to observe in raw electron microscopy micrographs^40^ (**Figure 5A**) and the length of these filaments is directly related to the reaction incubation time. Their growth kinetics have been studied in detail on both short and long time-scales^41–43^. Following a rate-limiting nucleation step by a cluster of 2-6 RecA monomers, filaments are rapidly extended by first-order reaction kinetics by sequential monomer addition^43^. In the presence of ATPγs, saturation of the oligonucleotides with RecA results in juxta-position of neighbouring filaments, resulting in concatenation^44–46^ and the appearance of longer filaments than would be expected from the contour length of a single oligonucleotide fully decorated with RecA. Moreover, the length of filaments on cryo-electron micrographs is easily quantifiable with basic image analysis tools.

**Figure 5.**
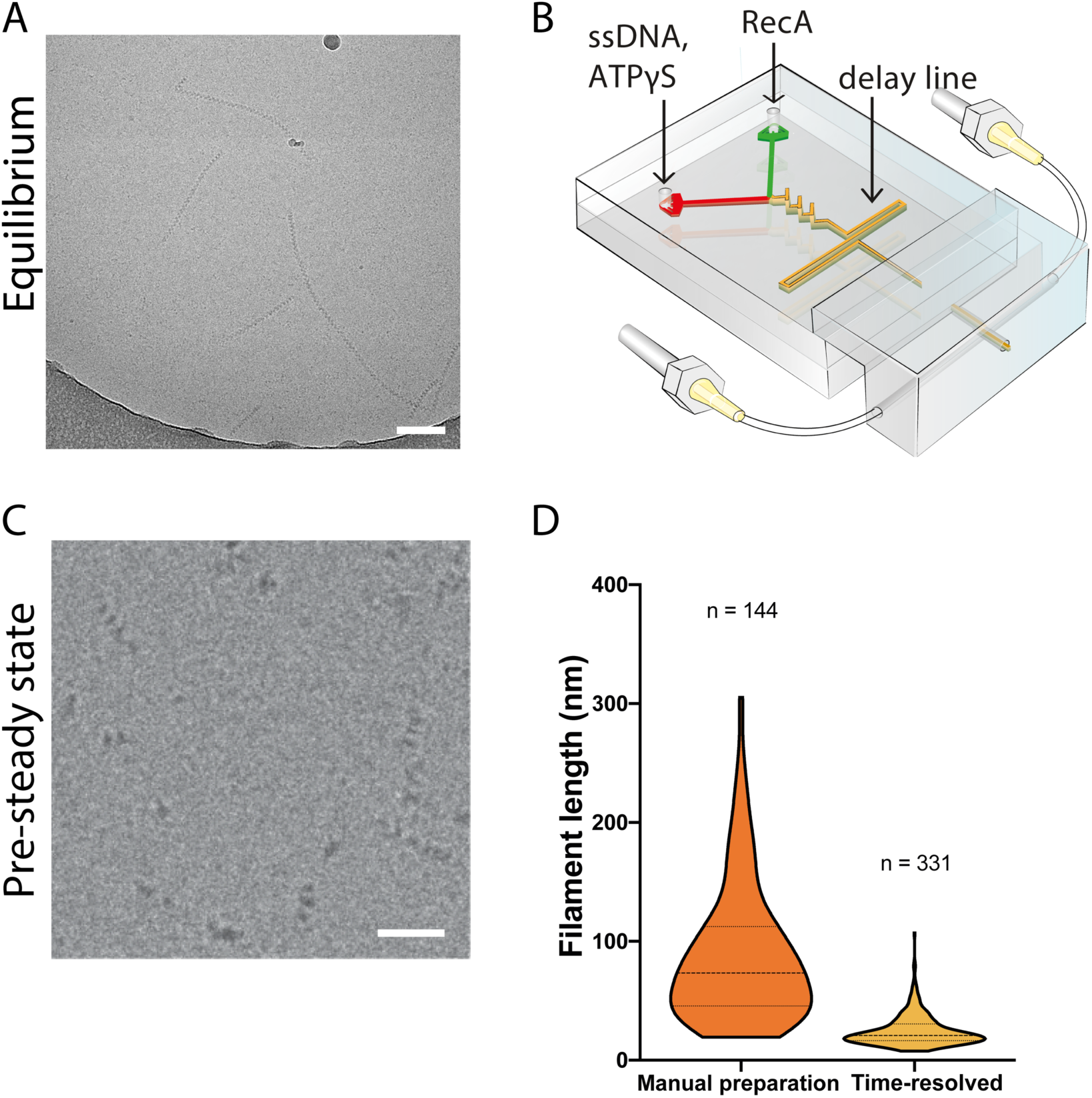
Observation of RecA-ssDNA filament growth by cryo-EM. **A** RecA filaments on a manually prepared grid. Scale bar is 100 nm. **B** Diagram of a mixing chip with a delay line. **C** RecA filaments observed on a grid prepared by blot-free spray-plunging in a chip with a 20 ms delay line. Scale bar is 50 nm. **D** Violin plot of quantified filament lengths from **(A)** and **(C)**. Dashed line indicates the median filament length, and the shape of the violin curve indicates proportion of filaments at different lengths.

To evaluate the ability to kinetically trap and image biochemical pre-steady state intermediates, as opposed to steady-state equilibrium structures, we performed parallel experiments. To this end, RecA was mixed with single-stranded DNA (ssDNA) and ATPγS, either manually or inside of a microfluidic mixing chip (**Figure 5B**), followed by cryo-EM grid preparation. For the manual mixing and grid preparation, the incubation was several minutes and resulted in an equilibrium of different filament lengths (**Figure 5A, D**). For the reaction incubation inside a chip we adapted the microfluidic concept that path length inside the device determines the incubation time, provided that velocity of the fluid is tightly controlled^47^. Thus, we introduced a delay line downstream of the mixing region, resulting in a nominal in-chip incubation time of around 20 ms (**Figure 5C, D, Supp Figure 5A, B**). We observed nucleated RecA filaments at the earliest time points of filament formation. As expected, observed RecA filament lengths were much shorter (**Figure 5C**), and the length distribution was markedly reduced compared to the manual preparations (**Figure 5D**). The asynchrony in lengths following manual preparation is easily understandable due to the fact that observed filaments have reached the equilibrium of provided reactants and are no longer undergoing first-order filament extension.

The deviation from the mean in the microfluidic experiments, albeit much smaller, merits further attention. One source of experimental inaccuracy in microfluidics may arise from the interaction of the reactants with the surfaces of the chip. To measure this effect, we quantified the adsorption of protein, DNA, and ATP following passage through PDMS microfluidic channels^48^ (**Supp Figure 5C**). We observed differential interactions of the reactants, where 19% of protein, 29% of DNA, and 42% of ATP were adsorbed by PDMS during a one-second incubation. Variance in adsorption is likely due to different diffusivity and electrostatic properties, which are expected to vary slightly depending on the reactants used.^49^ Nevertheless, these values would not result in rate-limiting depletion of the reactants. The slow nucleation of RecA filaments generates some overall error in reaction initiation as a proportion of filaments will only start growing after they have travelled an uncertain length along the delay line. Thus many filaments of shorter length than the average likely originated from such late nucleation events.

In contrast, the outliers towards larger than average length filaments likely arise due to the fluid dynamics of the system. The Reynolds numbers inside the used microfluidic chips under steady-state flow conditions are in the range of 100, resulting in laminar flow throughout the microfluidic channels, with the exception of the mixing region. This results in slower flow rates closer to the edges of the channel. Thus, the in-chip residence time, equivalent to incubation in a biochemical reaction, is characterised by a non-Gaussian distribution instead of a sharp peak which is, on average, equal to the nominal time but tends to be significantly longer for a subpopulation of reactions. Indeed, computational fluid dynamics simulations (**Supp Figure 5D**) and particle tracing predicted a characteristic residence time distribution as expected by the Taylor dispersion in laminar flow^50,51^ (**Supp Figure 5E**). These results are in good overall agreement with the observed filament length distribution (**Figure 5D**) and suggest that errors due to residence time scattering in the microfluidic chip are likely to be the dominant source of inaccuracies in time-resolution experiments conducted with this or similar systems.

### Pre-steady state RecA-ssDNA filament growth kinetics determined by trEM

Finally, having thoroughly characterised the scope of trEM operational parameters and their limitations, we designed and performed a proof-of-concept experiment aimed at obtaining kinetic information solely through image analysis of pre-steady state structural changes visualised by trEM. We performed a time-course of RecA filament growth reactions on sub-second timescales spanning three orders of magnitude to determine the fidelity of final time-resolution. This was done by constructing a series of microfluidic devices with varying lengths of the incubation channel that defines specific reaction times (**Supp Figure 6A**). In each chip, purified RecA was mixed with ssDNA and ATPyS, and the reactions sprayed onto EM grids. Each timepoint of RecA filament growth was captured in at least three independent experiments, on separately prepared reaction mixtures on different days to test for the reproducibility of the method. The length of RecA filaments was then measured at each time point by cryo-EM analysis with, on average, several hundreds of individual filaments per replicate (**Figure 6A, B**). By determining median filament length per replicate, we obtained linear growth rates for RecA-ssDNA filaments of roughly ∼20 nt/s, which closely matches previously reported growth rates observed over orders of magnitude longer time scales^41^. This was done by assuming that each RecA monomer holds three DNA bases and extends the filament by 1.53 nm^41^. This showed that we could robustly resolve pre-steady state growth of RecA filaments at the earliest timepoints using only structural information as a readout (**Figure 6A, Supp Figure 6B**). Moreover, there was little experimental variation for each time point (**Supp Figure 6D**). These results quantify for the first time the fidelity of time-resolution achievable by trEM, and demonstrate that the developed time-resolved cryo-EM method is robust and broadly applicable.

**Figure 6.**
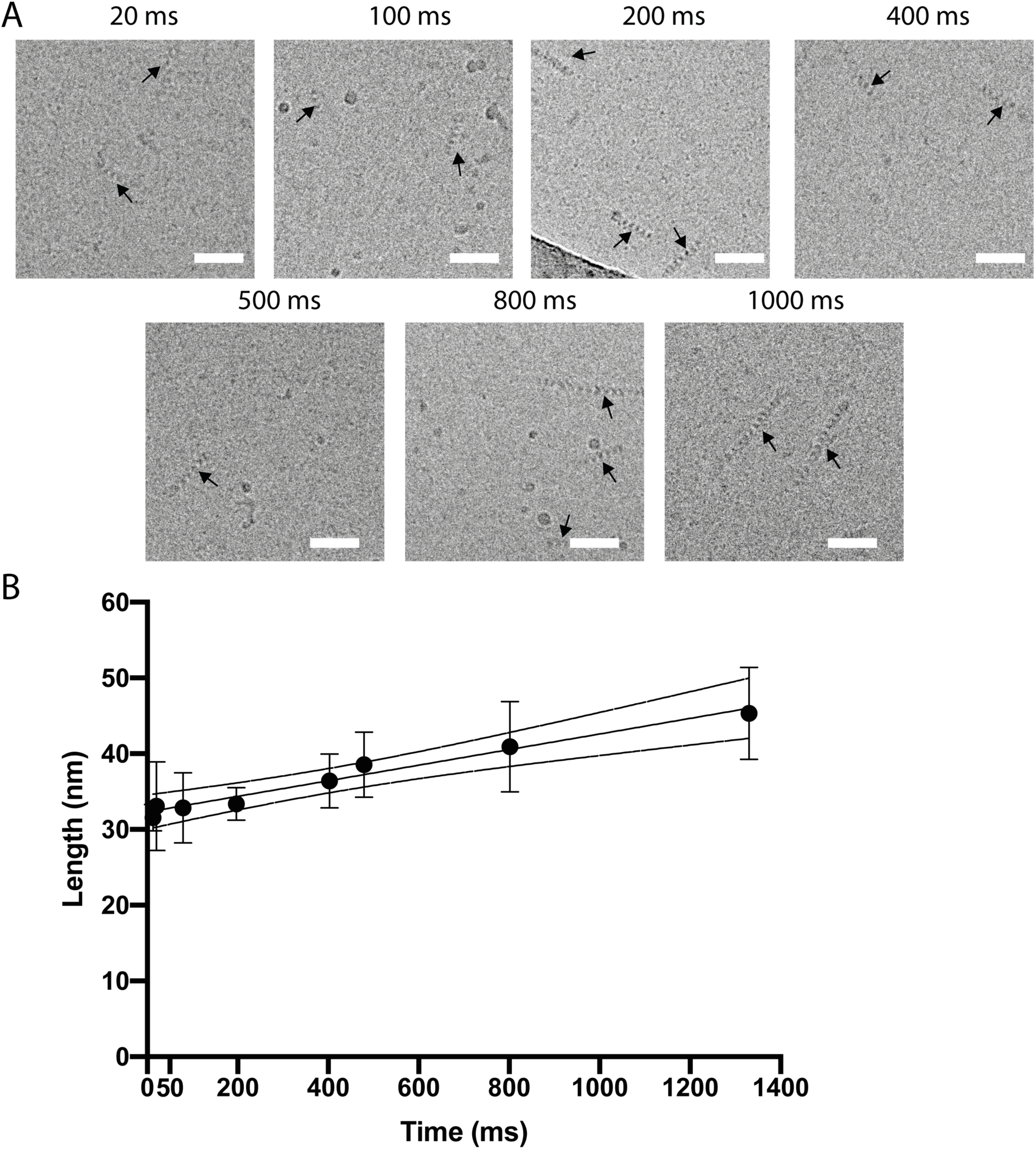
Kinetic analysis of RecA-ssDNA filament growth by time-resolved cryo-EM. **A** Representative micrograph of each studied time point indicated in milliseconds. Scale bars are 50 nm. **B** Growth curve quantifying the averages of the median lengths of at least three replicates per time point, shown are standard deviations (error bars) and the 95% confidence interval of the linear regression fit. R-squared of the fit through the means of medians is 0.973.

## Discussion

Structural biology provides highly detailed 3D models of biological macromolecular assemblies. However, the dynamic conformational changes required to perform specific functions often remain unobserved, emphasising the need to develop methods that can time-resolve biochemical reactions as they occur, without sacrificing molecular detail. Exploring such structural transitions, especially the intermediates that enable function, will provide a new window into understanding the very basics of biochemistry. In this study, we present a major step in the development of time-resolved structural studies, by developing a reliable and robust methodology for the initiation, synchronisation, and freeze-trapping of biochemical reactions onto cryo-EM sample support grids for subsequent analysis. Furthermore, we conducted a careful analysis of each step as it relates to the performance and limitations of the current technology and validated our method to quantify time-resolved growth of RecA-filaments. We obtained reliable biochemical growth rates for RecA-ssDNA filaments, on a par with biophysical growth rates determined over longer timescales^41,43,44^, demonstrating that trEM successfully combines biochemical and structural studies. Kinetic data obtained by trEM sample preparation and image analysis has the key advantage of not relying on fluorescent or radioactive labels. All reactants used in the microfluidic experiments were free in solution with no outside intervention or specific release mechanisms. This demonstrates for the first time that *de novo* biochemical reaction rates can be determined using fixation by vitrification of specific timepoints along the reaction path. A further major advantage of this technology is the ability to holistically study all structural transitions in the same experiment, rather than monitoring areas close to the chosen label sites, as done in routinely established biophysical methods. Future experiments will be designed to exploit these new opportunities and will undoubtedly greatly broaden the applications and biological insights obtainable by trEM.

The automated system for time-resolved cryo-EM sample preparation described here (**Figure 1B, Supp Figure 1A**) is versatile, simple to build and modify, and readily adaptable to different experimental needs. In particular, the microfluidic production pipeline ensures reproducibility in microfabrication and minimises cross-contamination by virtue of the very cost effective and simple replica molding of single-use PDMS microfluidic devices. The entire design and production process relies solely on affordable and readily available components and techniques, without the need to access clean room facilities, making it possible to implement this time-resolved system in most biological research institutions and thus widen its applications. Its two major features are a rapid blot-free sample vitrification method using a gas-assisted nozzle to generate a sample spray, and a tunable microfluidic mixing and incubation delay line, which can be readily customised to cover the vast majority of timescales relevant to biochemical processes.

The standard and most widely used method of cryo-EM sample preparation relies on manual sample application, followed by blotting of excess liquid and vitrification of the resultant thin film (**Figure 1A**)^28^. Although transformative for structural biology, this method also has limitations. For instance, the longer the sample spends in a thin film prior to vitrification, the likelier it is to contact the air-water interfaces potentially leading to denaturation^34^ or to adopt a preferred orientation, resulting in loss of structural information due to lack of viewing angles^52^. Recent improvements to the support grids and to how the sample is applied have begun to address these issues^33,34,53,54^. Here we describe a different approach, using standard grids rapidly plunging through a sprayed, atomised sample (**Figure 1B, Supp Movie 1**), faster than 30 ms. We determined a near-atomic resolution structure by single-particle analysis of an apoferritin sample prepared in this way (**Figure 4H, I** and **Supp Figure 4B**), which was practically indistinguishable from one obtained by standard preparation (**Supp Figure 4C**). However, the apoferritin particle distribution inside of the thin vitreous layers was markedly different between the two methods. Consistent with previous reports of rapid sample application onto self-wicking grids^35^, we demonstrated a close to completely uniform distribution for the rapidly prepared sample (**Figure 4F, G**), as opposed to more than half the particles being stuck to the interfaces following standard preparation (**Figure 4D, E**). Importantly, the sprayed sample spent less than 3 ms on the carbon grid before vitrification, compared to time spent using conventional methods. Future work will further clarify the relative contributions of blotting and preparation times. While apoferritin is not known to suffer from denaturation at the surface of the ice or from preferred orientations, this new method will likely facilitate structural studies of samples that are adversely affected by the air-water interfaces.

A second major limitation of the standard cryo-EM sample preparation method is that the process is too slow to capture intermediate states with lifetimes of milliseconds without the aid of mutagenesis, chemical inhibitors or crosslinking agents. To circumvent this limitation, we approached two further challenges. The first was rapid and complete mixing to initiate a biochemical reaction. Although diffusion equilibrates the concentrations of sample components over time, on average proteins diffuse less than a hundred micrometres per second, which prevents the synchronisation of biochemical reactants on sub-second time scales^55^. In the field of microfluidics, this problem has been successfully solved by the use of active or passive mixers which ensure complete mixing in milliseconds^56–58^. In this work, we incorporated a passive mixing geometry (**Figure 2A**) and experimentally determined the optimal combination of 3D mixing elements and flow rate to achieve near-complete mixing in about 3 ms (**Figure 2C**). Together with the blot-free sample application by spraying onto an accelerated plunging grid (**Figure 3B, F**), this defines the total dead time of our method to be less than 30 ms, which ensures its applicability to studying many biochemical processes.

Having effectively established rapid and reliable mixing to define the starting point of a reaction, the second challenge was assessing the synchronisation over the course of the experiment inside the microfluidic delay line (**Figure 5B**). Importantly, time-resolved growth of RecA filaments experimentally demonstrated significant enrichment of a characteristic length distribution at all studied time points (**Figure 6, Supp Figure 6B**), implying that the current setup allows a label- and fixation-free method to study transient intermediates. Optimal synchronization is governed by the residence time distribution (RTD) of the microfluidic geometry at the specific flow condition. We assessed the RTD under our experimental conditions by computational fluid dynamics (CFD) simulation analysis (**Supp Figure 5C, D**) and found that our simulations reliably predict experimental outcomes (**Figure 5C, D**). RTD arises physically from the laminar flow in microfluidic channels at steady state, where due to dead zones in the geometry the reactants can incubate for an indefinite time before exiting^59^. This can generate overlap in populations, potentially complicating subsequent analysis. Since the RTD comprises the biggest limitation to achievable time resolutions, it is worthwhile considering stop-flow systems and geometries that could reduce its effects. An alternative approach could be to accurately determine the extent of RTD through CFD simulations and experimental model systems and treat it as a linear response function of the system, attempting correction by deconvolution.

The major advances reported here as well as by others^21,26^ prompt further meaningful improvements to trEM. Currently, the amount of sample consumed per experiment is large in comparison to other sample preparation approaches^33,60,61^. Due to the use of a microfluidic mixing and incubating device, continuous flow is required, which in turn necessitates several seconds of flow before reaching a steady operating condition when the flow is generated by a syringe pump. Using injection valves to maintain steady flow without unnecessarily spraying sample could in principle reduce sample consumption to sub-microliter volumes. The analysis of the droplet sizes and speeds generated by our nozzle (**Figure 3C, D**) suggest a hitherto unexplored but promising additional improvement – the use of gas-dynamic virtual nozzles^62,63^. These nozzles are capable of producing droplet jets with similar parameters at much lower fluid flow rates, further reducing sample consumption.

Apart from the above engineering challenges of improving time-resolved sample preparation for cryo-EM, single-particle analysis of pre-steady state datasets harbours particular difficulties. Multiple structurally similar states may co-exist and it will be critical to detect them reliably in order to quantify their relative populations at any studied time point, thus establishing the order of molecular events. Unbiased algorithmic mining of conformational heterogeneity by emerging techniques will be of paramount importance^7,64–67^.

The newly developed experimental approach to preparing time-resolved samples for cryo-EM and single-particle analysis enabled us to reliably and reproducibly obtain near-atomic spatial and two-digit millisecond temporal resolution. Furthermore, we demonstrate the ability of trEM to closely follow biochemical reactions by using the model system of RecA filament growth, obtaining kinetic rates from raw micrographs that are in agreement with existing biophysical data. Thus, since most major difficulties in adapting cryo-EM to time-resolved studies have been circumvented and/or characterised, we are confident that time-resolved cryo-EM will offer unprecedented insights into biochemical reactions, including the sequence of reaction intermediates and pathway stochasticity.

## Data availability

EM maps are deposited in the EM Data Bank under EMD-10712 (trEM) and EMD-10714 (Vitrobot). Technical drawings and other data are available from the corresponding authors upon reasonable request.

## Author contributions

R.I.E., M.P., S.S.L. and C.S.L. designed research. M.E.M, B.L., M.M.N.E, S.G.J. and S.S.L. designed and produced microfluidic devices and characterised mixing efficiency. D.S., B.L., S.S.L., M.E.M, M.M.N.E. and R.I.E. designed, produced and optimised the mechanical and electrical integration of the system. S.S.L., M.E.M. and R.I.E. designed, produced and optimised the nozzle and collected and analysed high-speed camera data. M.E.M., A.N. and R.I.E. collected cryo-EM data and performed data analysis. M.E.M, S.S.L and R.I.E performed and analysed CFD simulations. M.E.M. and R.I.E wrote the manuscript with contributions and input from all authors.

## Acknowledgements

We thank D. Patel for technical assistance in fabrication and microfluidic chip production.. N. Koglin designed and manufactured the electronics board. We are indebted to P. Rosenthal, R. Danev, D. Boehringer, I. Schlichting, B. Doak, M. L. Gruenbein, M. Pilhofer and J. Medeiros for discussions, help, and advice. We thank the ETH core facility ScopeM (P. Tittmann) and FIRST for support with imaging and microfabrication. We equally thank the Structural Biology (P. Walker, A. Purkiss), Advanced Light Microscopy (R. D’Antuono) and Making Lab (R. Desai, O. Z. Natali, C. Dix) scientific technology platforms at the Francis Crick Institute. Research in the Peter laboratory and the Lee laboratory were funded by the Global Research Laboratory (NRF-2015K1A1A2033054) through the National Research Foundation of Korea (NRF). In addition, Research in the Peter laboratory was funded by the Swiss National Science Foundation and the European Research Council (ERC). Work in the Enchev laboratory was supported by the Francis Crick Institute, which receives its core funding from Cancer Research UK (FC001597), the UK Medical Research Council (FC001597) and the Wellcome Trust (FC001597). The authors declare no competing financial interests.

## Methods

### Manufacturing of a setup for cryo-EM grid preparation by spray-plunge-freezing

The frame of the setup was manufactured from stainless steel in a standard mechanical workshop. A transparent plastic box encasing the entire set up was also custom made and connected to a humidifier (UHW 60065, Medisana). Modified N7 tweezers (Dumont) were attached through a custom-made mount to a servo motor (HT750, Highest Korea). The fluid pump was a syringe pump (PHD Ultra, Harvard Apparatus). 250 μl gas-tight glass syringes (1700 PTFE Luer Lock, Hamilton) were used with the pump. Arduino mega 2560 (Arduino) controls the position of the tweezer in the rotation arm, on/off of the spraying gas valve, and the humidifier. Positioning between spraying nozzle and tweezers was done with XYZ stage (BO9-47, Suruga Seiki). The grid was plunge-frozen in a standard vitrobot ethane/nitrogen container (ThermoFischer – part no. FEI0815S).

### Electrical board and software integration/control

The electronics board (**Supp Figure 1B, C**) was manufactured to apply correct voltage for both power and control to each component, as well as containing an option for a second servo, with voltage control between 7.5 and 5 V. Software control of the system was achieved by communication between the open-source electronic prototyping platform (Arduino) and a PC. The Arduino was running the Firmata library, allowing other PC software to relay commands to the Arduino. The actual control software was a simple user-friendly GUI written in Processing, using the controlP5 library for visual control elements. The pump was controlled directly from Processing using a serial control library.

### High-speed camera imaging and data analysis

High-speed camera imaging was done with Fastcam Mini AX200 (**Figure 3E, F**) or Nova S12 type 1000K **(Figure 3B, Supp Movie 1, 2)** (Photron) and a LED light source (F5100, Photonic) and quantified using VisiSize (Oxford Lasers). Lenses used were a 12x lens with fixed focus, 2x adapter, 0.5x and 2x attachment lenses (Navitar). Camera control was achieved by the manufacturer’s software (PFV4, Photron). LED light was directly behind the imaged area for optimal illumination, when needed. Standard imaging settings for sprays were 100,000 frames per second, 0.2 μs shutter speed, 1024×128 pixel resolution **(Figure 3B)**. For droplet landing imaging 80,000 frames per second, 0.2 μs shutter speed, 384×304 resolution **(Supp. Movie 1, 2)**. For imaging tweezers plunging, we used 10000 frames per second, 33.3 μs shutter (due to inability to position light directly behind), 1024×1024 resolution **(Figure 3E, F)**. Movies were processed by applying software normalization and pixel gain in PFV software prior to imaging. Postprocessing with High-Dynamic Range (HDR) was used for grid landing movies to compensate for the obstructed illumination caused by the grid passing between the camera and the illumination source.

### PDMS chip design and manufacturing

The nominal fluid velocity and therefore channel geometry per timepoint was calculated using the Poiseuille formula for frictional pressure drop in laminar flow, adapted for rectangular channels. For this purpose, a publicly available web-based microfluidic calculator (Dolomite Microfluidics) was used. Times were rounded to the nearest 10 ms in **Figure 6B, Supp Figure 6B**.

Wafers for all types of microfluidic devices were manufactured using photolithography with UV exposure on negative photoresist via UV aligner (MA8, Suss MicroTec) or a direct photolithography machine (MicroWriter ML3, Durham Magneto Optics) on 3” silicon wafers (PI-KEM - WAFER-SILI-0004W25). SU-8 2035 photoresist (Microchem) was spin-coated onto the wafer at 1055 rpm for 30 s to reach a desired thickness of 100 μm (± 3 μm). All wafers were quality controlled by optical depth profiling using the optical profiler using interferometry with a 10x objective (S neox, Sensofar Metrology). Chips were designed such that 3D mixing features were formed by alignment of the top and bottom layers of the PDMS channel. For replica molding, the conventional soft lithography technique with polydimethylsiloxane (PDMS, Sylgard 184, Dow Corning) mixture was applied. The PDMS was mixed 10:1 base to curing agent ratio, mixed for 10 minutes, degassed under vacuum, and poured onto the wafer. After hardening by baking at 110 degrees, the chips were cut out and bonded after plasma treatment for 30 seconds. After plasma activation both sides of the chip were aligned and pressed together under a microscope by hand. After baking the bonded chips overnight at 80 degrees, fused silica tubing (TSP 100375, BGB) was inserted and glued with transparent silicone glue (Elastosil E43, Wacker) at the outlet to form the exit tubing, which inserts into the nozzle.

### RecA filament growth reactions

RecA filament growth buffer contained 25 mM Tris-HCl (pH 7.5), 10 mM Magnesium-Acetate, 100 mM Sodium-Acetate, 1 mM DTT. Final reaction mixture also contained 1 mM ATP-γ-S, 10.57 μM RecA (0.4 mg/ml), 31.71 μM ssDNA (3-fold of RecA). The ssDNA used was 144-mer (TTTTTTCTCACACTCATTTTTTTTCTCACACTCATTTTTTTTCTCACACTCATTTTTTTTCTCACACTCATT TTTCCTATATTTATTCCTTTTCCTATATTTATTCCTTTTCCTATATTTATTCCTTTTCCTATATTTATTCCT). Reactions were performed at room temperature. For filament growth experiments, two different solutions were prepared. One contained buffer, ATP-γ-S and ssDNA the other contained buffer and RecA. These were loaded into glass syringes and then inserted in the microfluidic chip through silica tubing (TSP 100375, BGB) to initiate the reaction. All concentrations stated are final concentrations after mixing inside the device. Quantifications of filament lengths were done in FIJI software ^68^, by manually tracing filament lengths.

### Mixing quantification by fluorescence microscopy

Tile scans of the entire mixing channel were acquired using a Leica SP8 confocal microscope (pin hole at 1AU) over 300 μm height (61 z-slices, every 5 μm), using a 10X/0.4 objective (Leica HC PL APO CS2) with final pixel size being 1.0111 μm. Fluorophores used were FITC and Rhodamine 6G, excited with 488 nm and 561 nm laser lines respectively in sequential mode to minimize the cross-talk. Before imaging, the fluorophores flowed in the channel for 30 s to reach a steady-state flow. After every junction, the z-stack was resliced along the z-axis to obtain transversal cross sections images (re-slice made with FIJI software^68^) (**Figure 2B**). We analysed a region of 10 x 10 pixels at the centre of the channel to avoid artefacts from the channel walls. Quantification of mixing efficiency was done by comparing the cross section at the start of the channel to the ones after each junction, measuring the mixing index in terms of standard deviation of pixel values, using equations previously described^68^.

### Cryo-EM grid preparation

Quantifoil R2/2, holey, 300 mesh grids were glow discharged for 40 sec at 40 mA with a K100X glow discharger (EMS). For freezing a Vitrobot Mark IV (FEI ThermoFisher) was used at room temperature and 100% humidity. 4 μl of apoferritin (diluted to 2 mg/ml in PBSA, pH 7.4) (Sigma - A3641) was pipetted onto the grid, blotted for 3 seconds, and plunge-frozen.

### Time-resolved cryo-EM grid preparation

Grids were glow discharged for 40 seconds at 40mA using a K100X glow discharge device (EMS). Grids were held at a distance of 8 mm from the nozzle. Sample was sprayed at 666 μl/min of sample flow and 0.8 bar nitrogen gas. Plunging speed was roughly 1.1 m/s. Grids used in experiments: Quantifoil R2/2, 2 nm Carbon, 400 mesh (**Figure 4B**). Quantifoil R2/2, holey, 300 mesh (**Figure 4A**, high-resolution datasets). Lacey carbon film with ultrathin carbon 400 mesh (Agar Scientific) (**Figure 4C**).

### Single-particle analysis

High-resolution cryo-EM data were acquired on Titan Krios operated at 300 kV with a FalconIII detector in counting mode with 20 frames per movie and nominal magnification of 96000x (0.845 Å pixel size). The electron dose was 33.6 e/Å^2^. For the sprayed apoferritin dataset the defocus range was −0.5 to −3.5 μm, while the vitrobot apoferritin dataset had a range of −0.5 to −2.5 μm. The larger defocus was due to the slightly thicker ice in sprayed samples. Movies were motion-corrected by using MotionCor2 with 5 by 5 patches^69^ and CTF estimation was done by CTFfind4 on dose-weighted micrographs^70^. All particle picking was done by crYOLO using a model trained in-house^71^. Subsequent image processing was performed in Scipion and RELION-3, using PDB 1AEW^72^, filtered to 40 Å as a reference^73,74^.

|  |  |  |
| --- | --- | --- |
| Dataset: | Sprayed apoferritin | Vitrobot apoferritin |
| Detector | Falcon III in counting mode |  |
| Nominal magnification | 96000X |  |
| Pixel size | 0.845 Å/px |  |
| Defocus range | -0.5 to -3.5 $\mu\text{m}$ | -0.5 to -2.5 $\mu\text{m}$ |
| Collected micrographs | 1747 | 843 |
| Processed micrographs | 1090 | 788 |
| Particles picked | 133833 | 146581 |
| Symmetry | octahedral |  |
| Box size | 250x250 |  |
| 2D classification | 65202 particles after 3 rounds of 2D classification | 99271 particles after 2 rounds of 2D classification |
| 3D classification | 9804 particles after 1 | 19485 particles after 1 |

|  | round of 3D classification | round of 3D classification |
| --- | --- | --- |
| FSC Threshhold | 0.143 | 0.143 |
| Final resolution (masked) | 2.77 Å | 2.89 Å |
| EMData bank deposition | EMD-10712 | EMD-10714 |

### Tomography of particle distributions

Tilt series were collected in Tomography 4.0 on a Talos Arctica (Thermo-Fisher Scientific, Waltham, MA) operating at 200 kV and with a Falcon III direct electron detector. The tilt series were collected in linear mode with a range of ± 54 degrees and 3 degree increment at a pixel size of 5.18 Å/px. Each exposure received a dose of 3 e/Å^2^ for a total dose of 111 e/Å^2^. The defocus used was −8 μm. Tomograms were aligned in IMOD^75^ and reconstructed with weighted back-projection followed by median-filtering and down-sampling by a factor of four. Template-matching of tomograms was performed with MolMatch software^76^. A low-pass filtered apoferritin map at 20 Å resolution was used as a template that was systematically translated across the tomogram before calculating the normalized cross-correlation with the tomographic volume. This procedure yielded a composite map containing the maximum cross-correlation coefficients (CCC) for each voxel in the tomogram. The top ∼2000 CCCs and their 3D coordinates were extracted from the map using the AV3 toolbox in Matlab (av3_createmotl)^77^. After plotting the CCCs, extremely high (false positives) or low (false negatives) values were excluded with an empirically determined cut-off threshold. Any remaining false positives and negatives were manually pruned using the EM Package for Amira 5.3^78^ (Thermo-Fisher Scientific, Waltham, MA). The remaining ferritin matches were displayed in Amira 5.3 and their corresponding Z-coordinates were used for quantification. To facilitate comparison, we binned the z-height of each tomogram into 10-percentile bins and counted the total number of apoferritin particles located per bin across all tomograms to produce the plots and quantifications in **Figure 4E, G**.

### Simulations of residence time distribution

Simulations were performed using COMSOL Multiphysics 5.4. Laminar flow simulations were performed with 3D models of the microfluidic device’s geometry. These were generated in Solidworks Professional Research 2019 (Solid Solutions) and simulation meshes were subsequently generated by using COMSOL built-in mesh generation tools using the fluid-dynamics preset. The stationary solution of the flow profile was then used as input, described by the Navier-Stokes equation, into a time-dependent study of particle tracking in the geometry. Proteins were approximated by 10 nm solid spheres with the molecular weight of 10 RecA monomers (378 kDa). Both drag forces from the flowing liquid and Brownian motion of particles were modelled as part of RTD analysis. Quantification was done by storing the total residence time of each particle inside the geometry from the inlet, measured at the outlet. Graphs were generated in Prism 8 using the “high” level of violin plot smoothing.

**Supplementary Figure 1.**
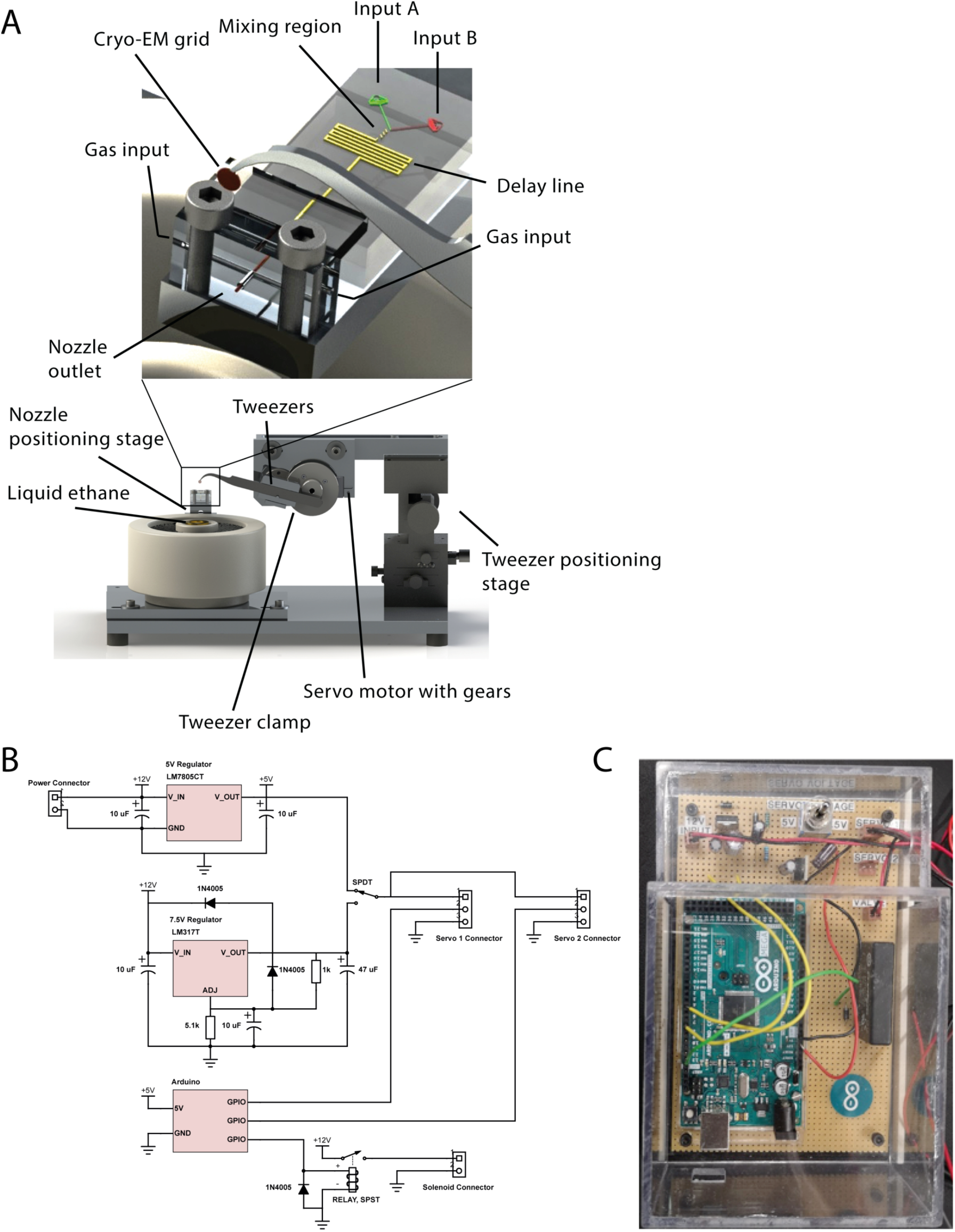
**A** Three-dimensional technical rendering of the set up for time-resolved cryo-EM sample preparation, indicating key elements. **B** Diagram of the electrical control board. It contains a voltage regulating circuit to manage power to different components as well as input/output from the Arduino. Not shown are the connections of the pump and camera, which connected directly via USB or ethernet to the Computer. **C** Image of the electronics board in a protective plastic case.

**Supplementary Figure 2.**
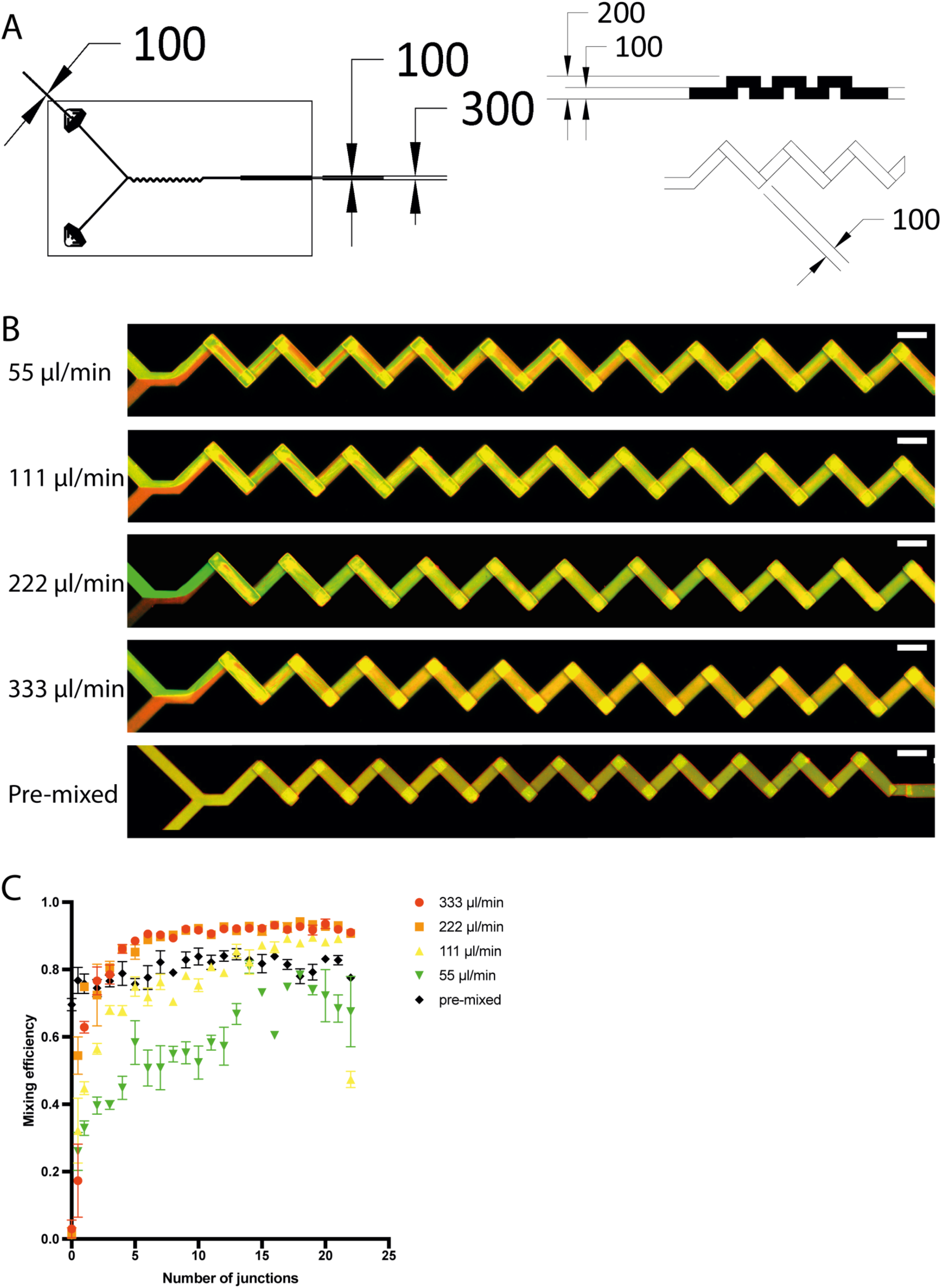
**A** Tcehnical drawing of the overall microfluidic geomertry and 3D passive mixing elements (left).Top view of overall microfluidic gemometry, (right top) side view of 3D passive mixing element, and (right bottom) top view of the 3D passive mixing element.dimensions in μm. **B** Confocal micrographs of mixing two fluorescent dyes at indicated steady state flow rates. Scales bars are 200 μm **C** Quantification of mixing efficiency data shown in **B** as a function of 3D mixing elements.

**Supplementary Figure 3.**
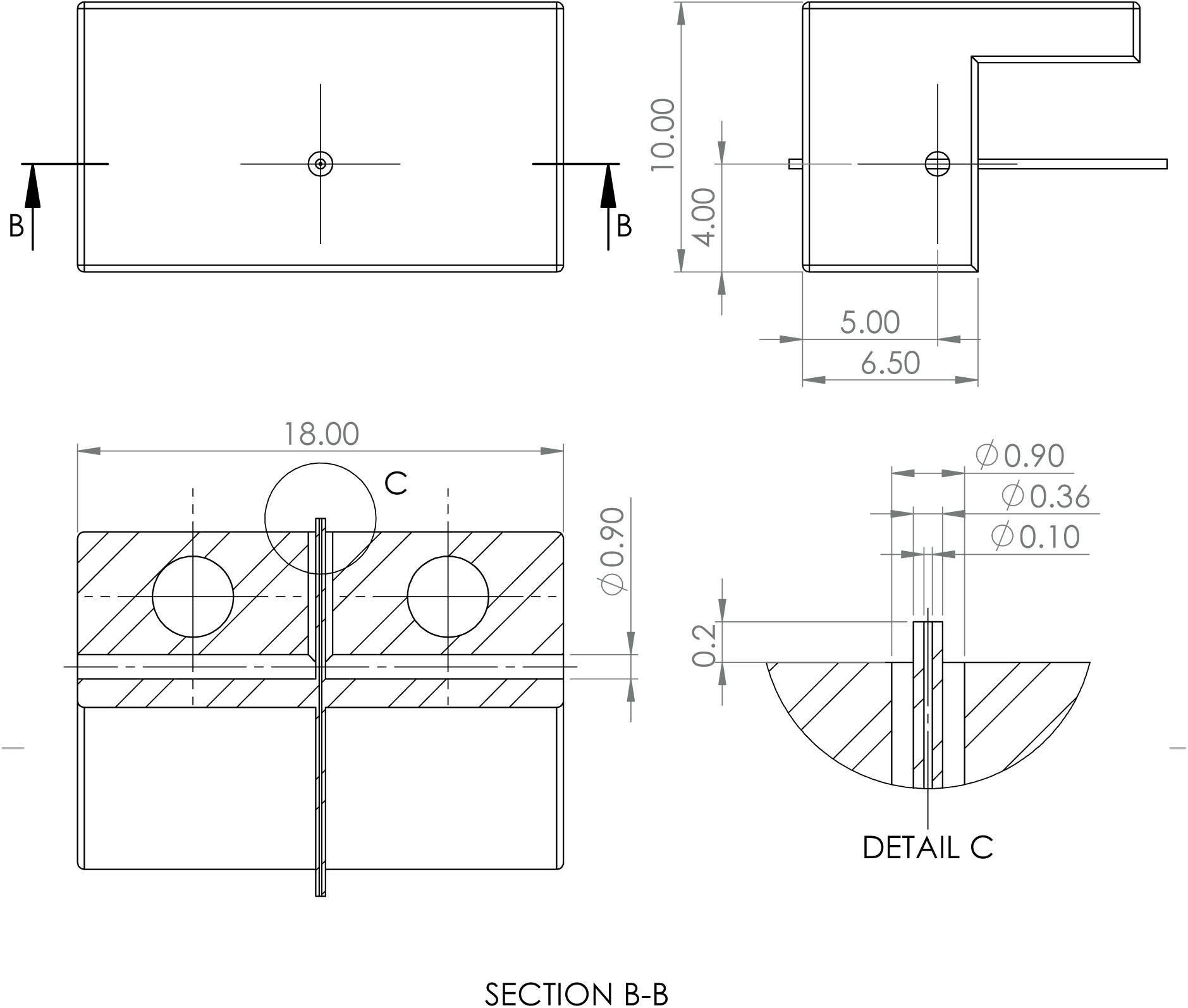
Technical drawing of the nozzle. Dimensions in mm.

**Supplementary Figure 4.**
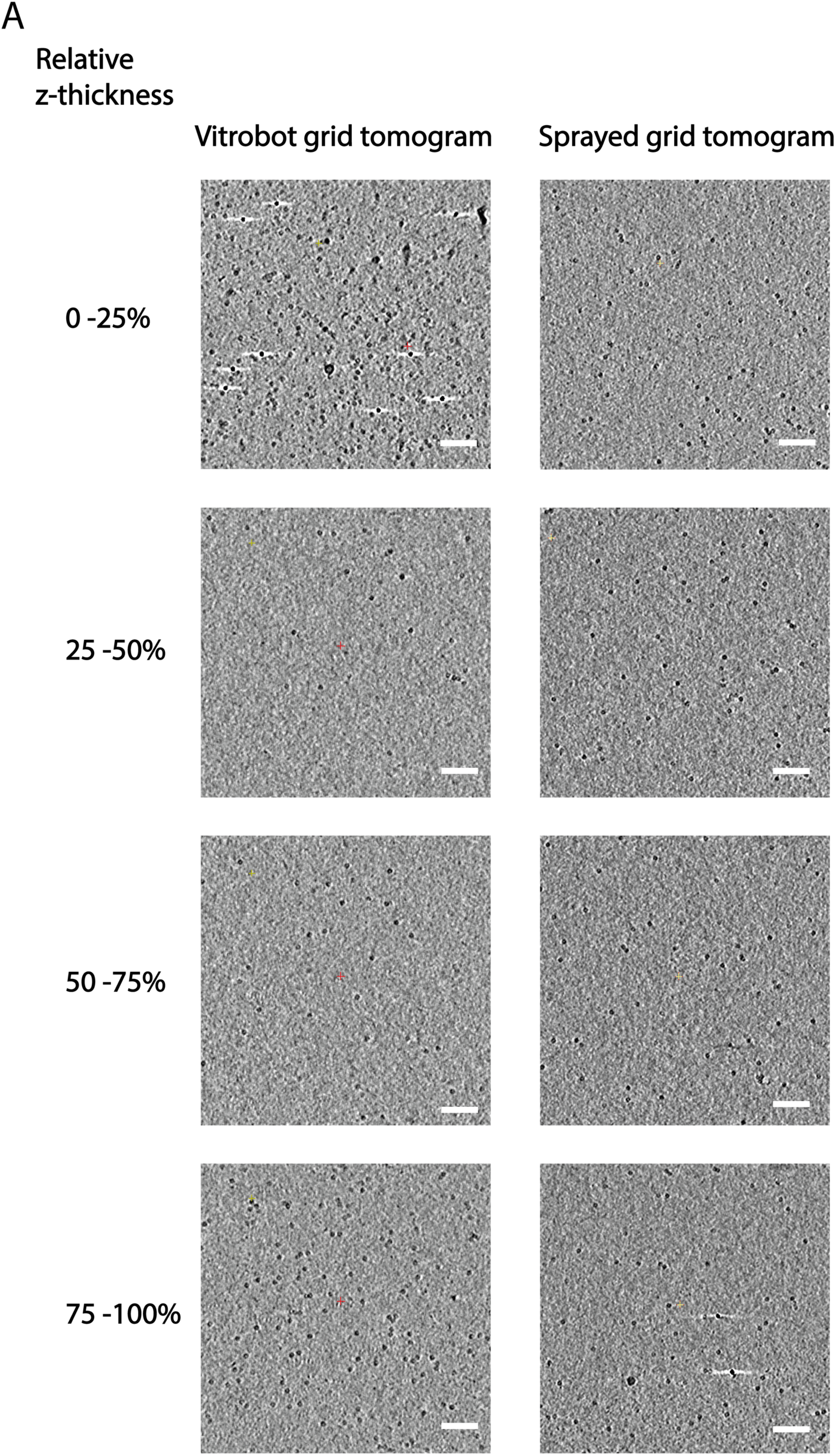
**A** Slices through tomographic reconstructions at indicated relative depths. Scale bars are 100 nm.

**Supplementary Figure 4.**
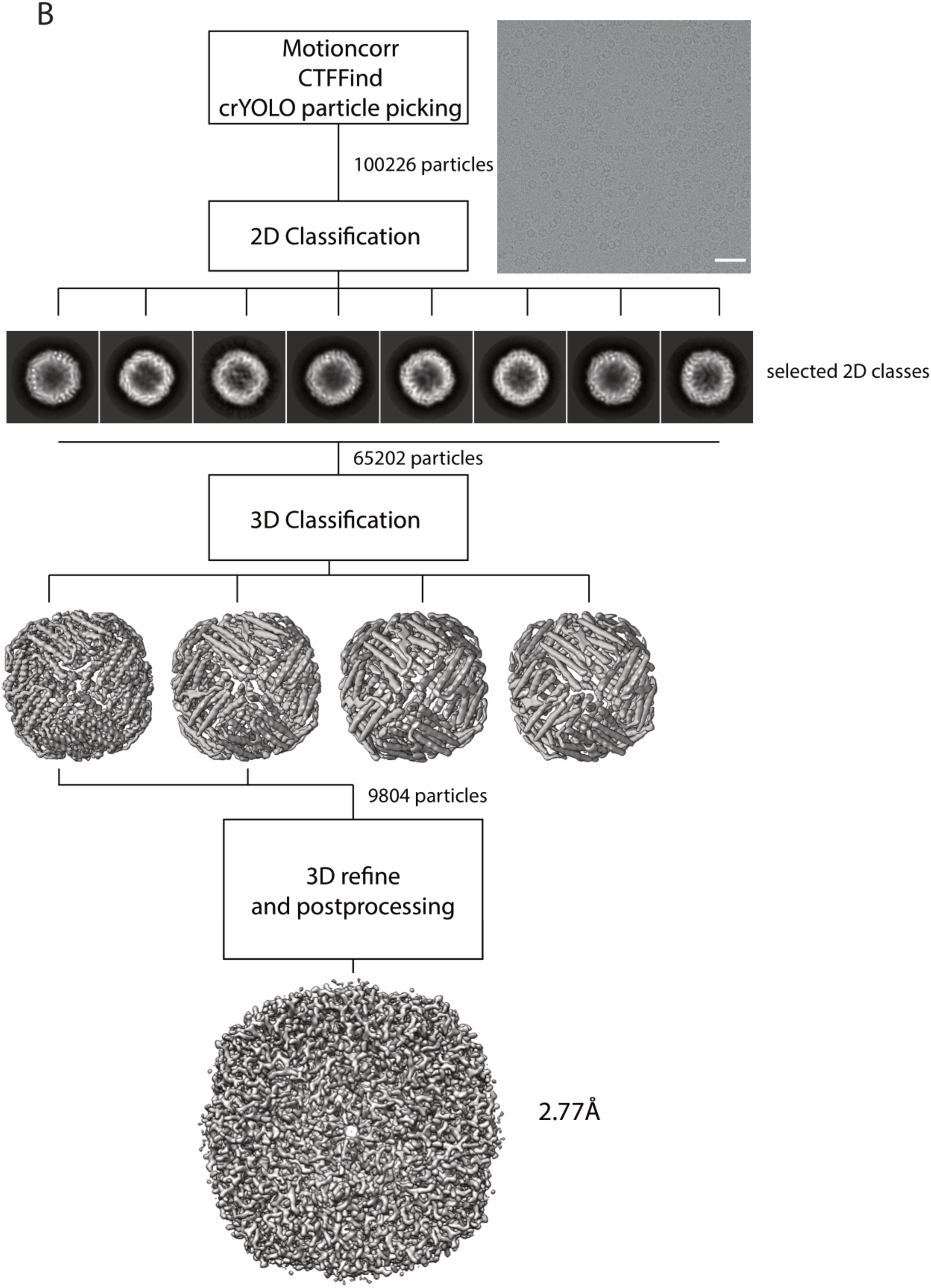
**B** Workflow of single-particle analysis and statistics of apoferritin from a sample prepared by blot-free spray-plunging. Scale bar 50 nm.

**Supplementary Figure 4.**
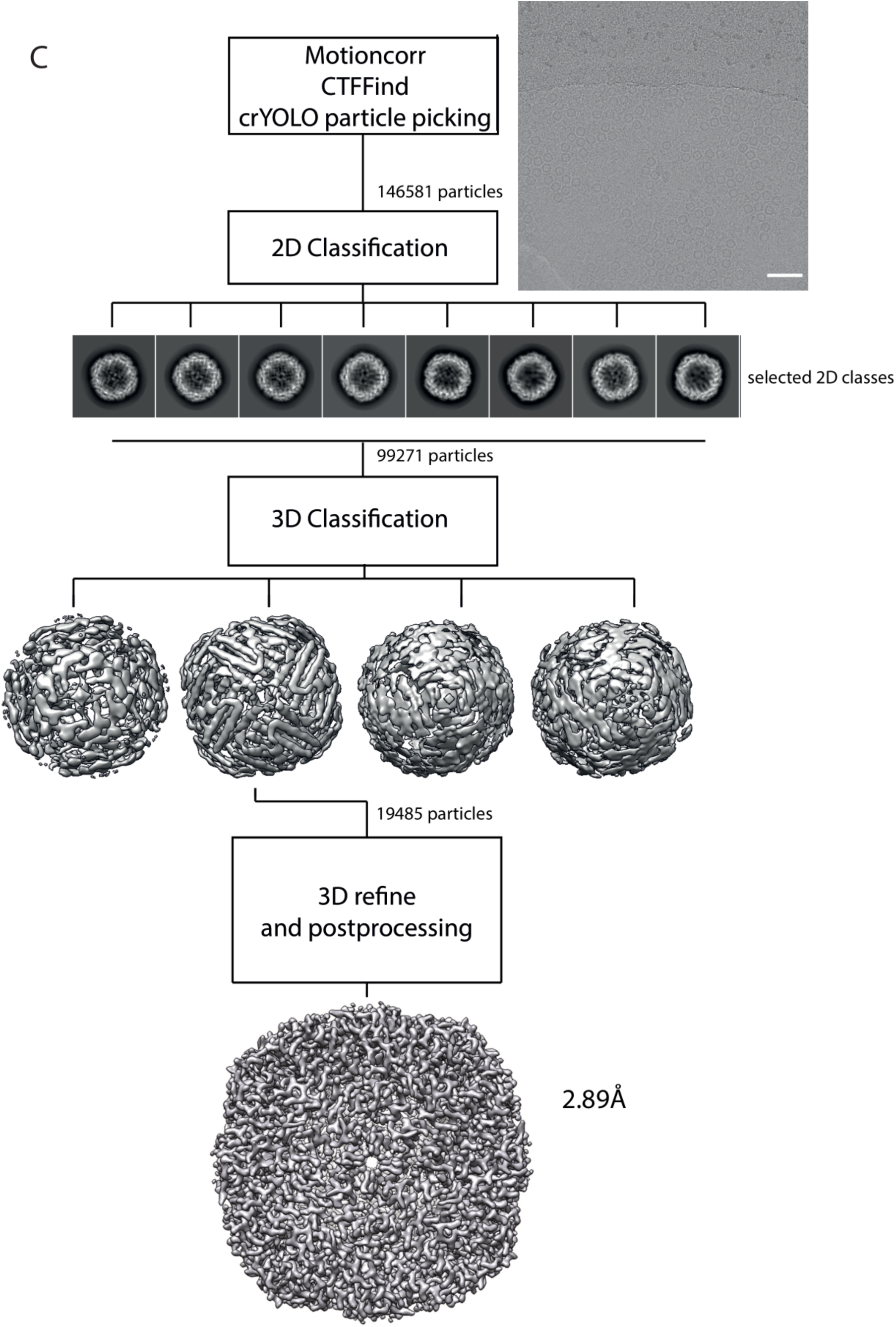
**C** Workflow of single-particle analysis and statistics of apoferritin from a sample prepared by Vitrobot. Scale bar 50 nm.

**Supplementary Figure 5.**
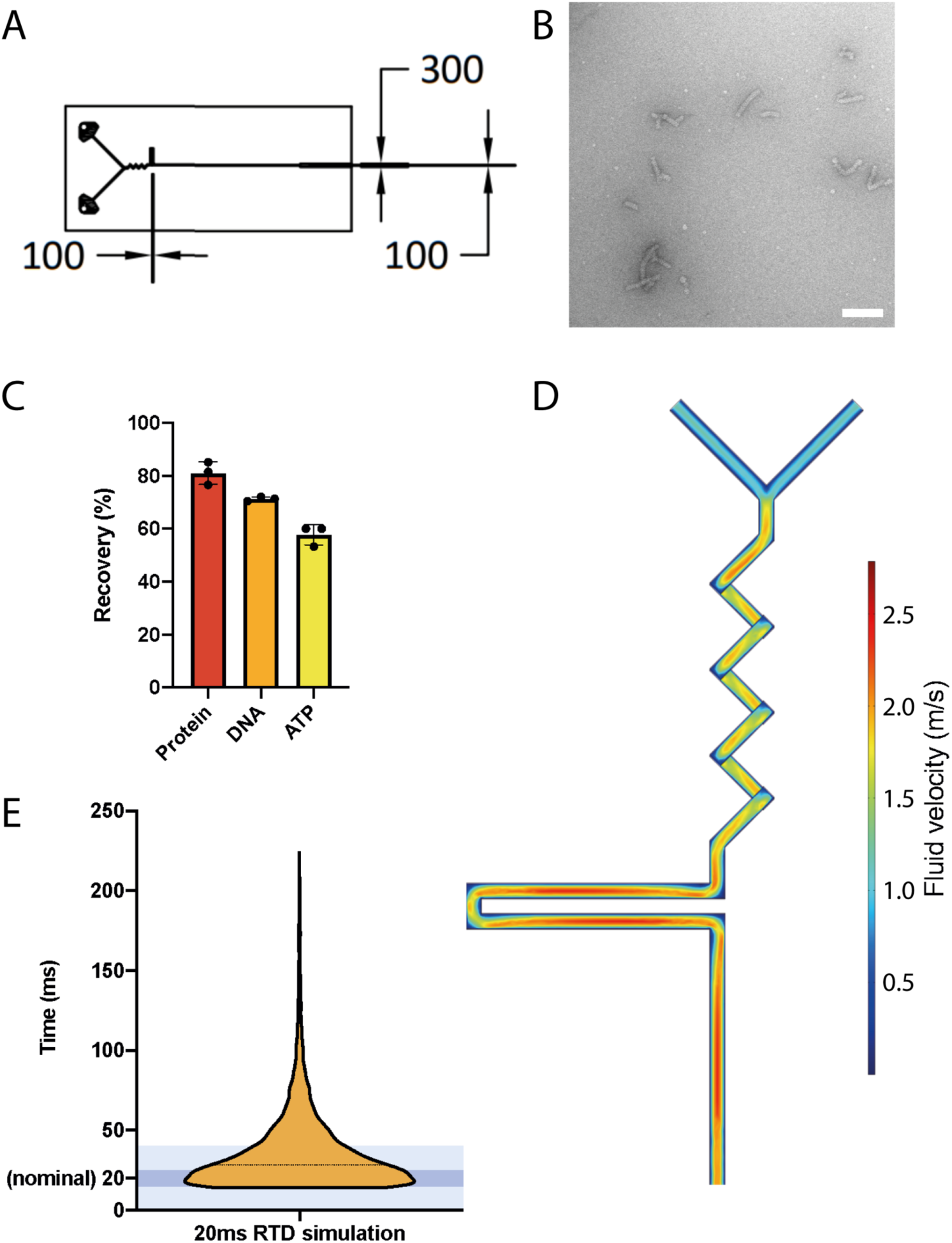
**A** Technical drawing of the chip with a 20 ms delay line used for **Figure 5C** (dimensions in μm). **B** Negative stain grid example produced using trEM. Negative stain can be used as a suitable alternative to freezing in ethane, as it can stop biological reactions in as little as 10 ms, with the caveat that most high-resolution conformational information is lost^81^. Conversely, it is well suited for initial proof of concept screening due its ease of use. The workflow for producing such samples was identical to the one described for cryo conditions, except for substituting the ethane cup for a custom made plastic cup which contains 500 μl of 2% uranyl acetate. **C** Quantification of sample loss to the PDMS chip. **D** Velocity of flow through the chip, which is computationally simulated using the conditions shown in **Figure 5C** CFD simulation of flow through the chip in **Figure 5C**. **E** Histogram of the simulated residence time distribution. Dark blue is 15-25 ms and contains 33% of total data. Light blue is 0-40 ms and contains 70% of total data.

**Supplementary Figure 6.**
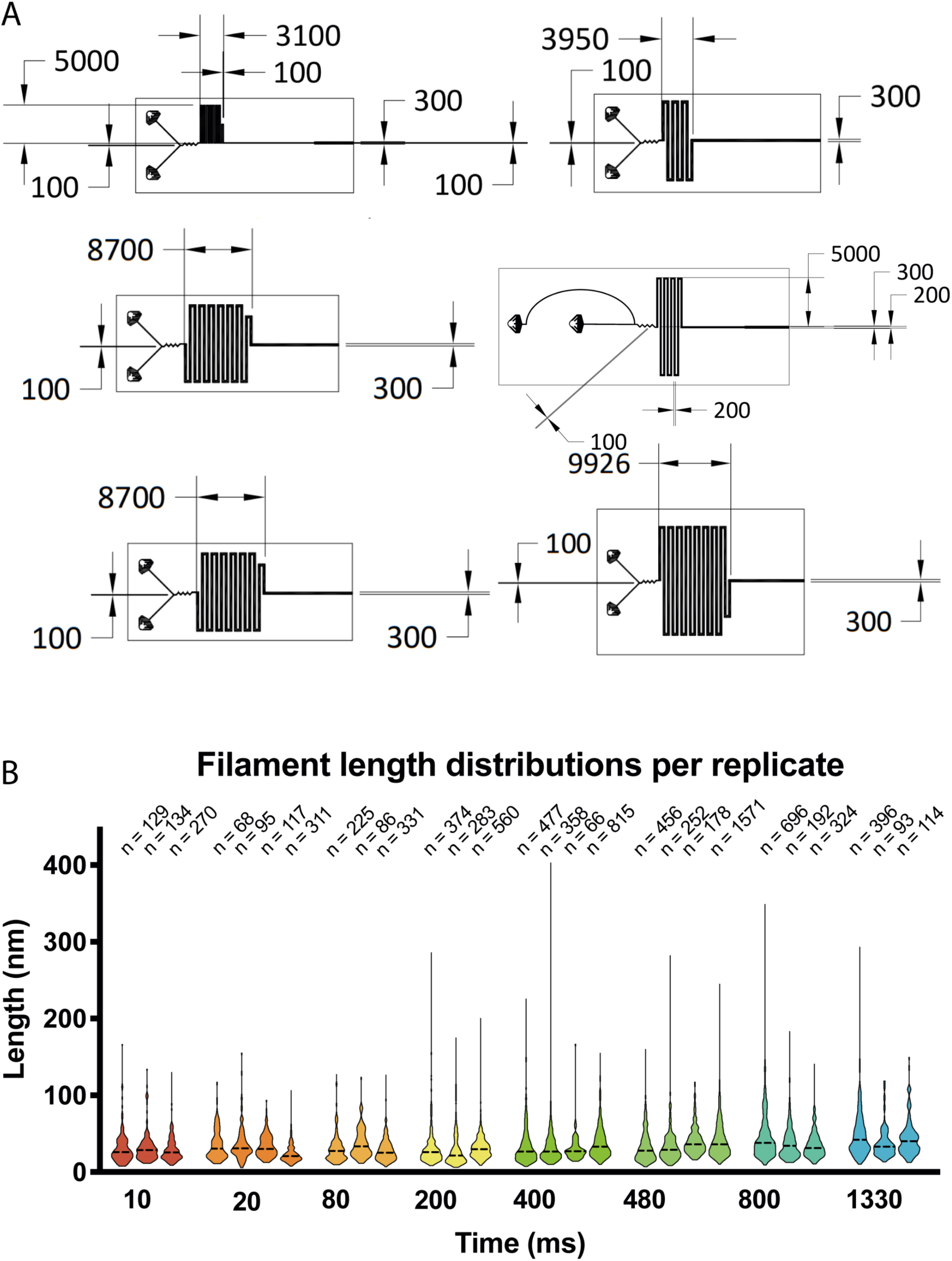
**A** Technical drawings of microfluidic chips and delay line designs used for producing the data in **Figure 6A** (dimensions in μm). Z-height of delay line 100 μm in first three panels, 200 μm in last three panels. **B** Violin plots of individual data points whose medians were averaged for the growth curve shown in **Figure 6B**.

